# Agency plans are inadequate to conserve US endangered species under climate change

**DOI:** 10.1101/647396

**Authors:** Aimee Delach, Astrid Caldas, Kiel Edson, Robb Krehbiel, Sarah Murray, Katie Theoharides, Lauren Vorhees, Jacob W. Malcom, Mark Salvo, Jennifer R. B. Miller

## Abstract

Despite widespread evidence of climate change as a threat to biodiversity, it is unclear whether government policies and agencies are adequately addressing this threat to species^1–4^. We evaluate species sensitivity, a component of climate change vulnerability, and whether climate change is discussed as a threat in planning for climate-related management action in official documents from 1973-2018 for all 459 US animals listed as endangered under the Endangered Species Act. We find that 99.8% of species are sensitive to one or more of eight sensitivity factors, but agencies consider climate change as a threat to only 64% of species and plan management actions for only 18% of species. Agencies are more likely to plan actions for species sensitive to more factors, but such planning has declined since 2016. Results highlight the gap between climate change sensitivity and the attention from agencies charged with conserving endangered species.

## Introduction

Climate change is a threat to ecosystems and biodiversity globally^5,6^, and has emerged as a driver of observed and potential species decline and extinction^7–9^. Government laws and policies should play a vital role in supporting climate change adaptation for imperiled species, yet imperiled species protections have been critiqued as insufficient in Australia^10,11^, Canada^12^, and Europe^13^. Funding shortfalls for environmental programs mean that governments may not be adequately addressing baseline threats to species, let alone more complex emerging threats from climate change^14,15^. Furthermore, the politicization of climate change in many countries, including the US, has led to different levels of concern and action on the topic among political parties^16,17^. Understanding whether and to what extent government authorities are supporting climate change adaptation, especially for imperiled species, is critical for improving tools and processes to reduce climate change impacts on biodiversity^18,19^.

The primary law directing the conservation of imperiled species in the US is the Endangered Species Act^20^ (hereafter, ESA). Central to the listing and recovery processes under the ESA is the enumeration and abatement of threats to species. The law directs the Secretaries of the Interior and Commerce to use the “best available scientific and commercial data” to make listing determinations on the basis of five threat factors: 1) habitat destruction and degradation, 2) overutilization, 3) disease or predation, 4) inadequacy of existing protections, or 5) other factors. While each factor may result from or be exacerbated by climate change, this threat is not explicitly described among the five factors. This is likely because the ESA was most recently amended legislatively in 1988^21^, the same year as the formation of the Intergovernmental Panel on Climate Change and four years before the first detailed discussion the consequences of climate change for biological diversity in the US^22^.

Nonetheless, the two agencies responsible for implementing the ESA, the US Fish and Wildlife Service (FWS) and the National Marine Fisheries Service (NMFS), have explicitly recognized the threat that climate change poses to species and the need to manage for its impacts. The FWS first described climate change as a threat in its January 2007 proposal to list the polar bear (*Ursus maritimus*) as threatened. Later that year, discussion of climate change appeared in recovery plans for the Indiana bat (*Myotis sodalis*) and Hawaiian monk seal (*Monachus schauinslandi*) and in five-year reviews for the red wolf (*Canis rufus*) and five sea turtle species (for references to species ESA documents, see archived data). The only assessment of climate change in ESA documents to date (to our knowledge) found that by the end of 2008, 87% of species recovery plans still did not address whether or not climate change was a threat^18^. The scientific community has identified climate change as the “primary threat” to nearly 40% of ESA-listed animals and over 50% of ESA-listed plants in the US^14^, and agency options for climate-related management action under the ESA have been available for over a decade^23^. Thus, it is vital to understand whether the lead agencies responsible for endangered species conservation have improved the use of their authority to help species adapt to the threat of climate change.

To determine if threats from climate change are being addressed by US agencies, we compared the climate change sensitivity of species to agencies’ discussion of climate change and plans for managing climate change threats for the 459 ESA-listed endangered animals found within US lands and waters. Because climate change sensitivity had not been systematically assessed for many of these species, we developed a trait-based climate change sensitivity assessment^24^. This assessment is a simplified version of existing tools (see Methods) and provides a preliminary evaluation of whether and which species’ life history and biological characteristics contribute to sensitivity to climate change (see Table 1). We focused on sensitivity (and related traits sometimes characterized as measures of adaptive capacity) because these, rather than exposure, are the elements of vulnerability that management plans can address^10^. Furthermore, because of the small populations and range sizes of many of the species we evaluated, available exposure tools may not accurately capture granular scale and stochastic effects in a meaningful way^25^. Focusing on sensitivity greatly reduced the time required to assess each species, allowing the assessment to be applicable to large groups of species, such as the >2,300 US and foreign species listed under the ESA. Our assessment relies on affirmative statements about relevant aspects of biology and life history; certain traits had to be identified in the literature for a species to be determined sensitive, and a species was considered not sensitive by default. Therefore, the assessment represents a conservative estimate of sensitivity and likely underestimated the actual sensitivity for some poorly-studied species or those for which that information was not described in publicly available sources.

**Table 1.** Questions in the rapid sensitivity assessment related to eight climate change sensitivity factors.

| Factor | Question and description |
| --- | --- |
| Temperature | Does the species have specialized thermal tolerance or depend on habitat with an important temperature threshold? Species were considered temperature sensitive if available information indicated the species has or depends on habitats with obligate or preferential temperature thresholds (e.g., sea ice). |
| Hydrology | Is the species dependent on habitat with a specialized hydrology? Species were considered sensitive if available information indicated they require narrow ranges of water depths, flow rates, timing, or seasonality (e.g., vernal pools or intermittent streams). |
| Disturbance | Is the species or its habitat sensitive to or dependent on a specific disturbance regime? This includes species in fire-adapted systems, species that rely on certain flood regimes, and species impaired by disturbance, such as old-growth forest obligates and species sensitive to excessive flooding. |
| Isolation | Is the species or its habitat geographically restricted or does it face intrinsic or extrinsic barriers to shifting its range to maintain its climate space? While many endangered species are found in small, isolated populations, we scored species in this category if available information indicated they are confined to mountains, islands, or headwaters; are narrowly endemic to spatially discrete habitats, like caves, springs or rare soil types; or if species movement to other suitable habitat is limited by habitat loss, development, dams, or other anthropogenic pressures. |
| Injurious species | Is the species or its habitat threatened by an invasive species, pest and/or disease organism that might benefit from climate change? We did not consider the species in question sensitive where the injurious species is ubiquitous or human-oriented (e.g., cats, rats, livestock). |
| Chemistry | Is the species sensitive to changes in chemical concentration, such as atmospheric CO<sub>2</sub>, water pH, or dissolved oxygen? |
| Phenology | Does the species rely on specific triggers for life cycle events, such as breeding, migration, or color change, that are likely to become out of sync with seasonal changes in resource availability or environmental conditions (i.e., phenologic mismatch)? |
| Obligate relationships | Is the species dependent on one or a few species such as a host, dominant food source, with limited alternatives if the required species declines due to climate change? We did not consider the species sensitive if it requires a host but can succeed in association with four or more species. |

After assessing species sensitivity, we determined whether climate change was described as a threat for species by reviewing official ESA documents published by FWS and NMFS. All endangered species have listing determinations, and most have either critical habitat designations, five-year reviews, recovery plans, or recovery outlines. We focused on the most recently published one or two of these types of documents to determine if climate change was described as a threat. We then determined whether these agencies planned management action to address climate change threats as part of species recovery by evaluating the same ESA documents (excluding species whose only ESA document was a listing decision, as these are not management-oriented). We tested if species sensitivity was a significant predictor of whether species ESA documents contained discussion of climate change as a threat and to what extent federal agencies planned to respond to climate change impacts. Data and results of the study are available in a free, interactive web application at https://defenders-cci.org/app/ESA_climate/.

## Results and Discussion

We found that nearly all endangered animals are sensitive to climate change impacts. All but one (Hawaiian goose [*Branta sandvicensis*]) of the 459 species (99.8%) are sensitive to at least one of the eight sensitivity factors (Table 1), and three-fourths (74%) are sensitive to three or more factors (Fig. 1a). However, agencies describe climate change threats in documents for only slightly more than half of species we assessed (64%; Fig. 1b) and plan management actions to address those threats for only a small fraction of species (18%; Fig. 1c). Logistic regression indicated that the number of sensitivity factors is a strong predictor of whether ESA documents discussed management action (F(1,419)=6.57, β=−0.31, p<0.01; Fig. 1a). Agencies are more likely to plan climate adaptation management actions for species that are sensitive to more climate factors than for species that are sensitive to fewer factors; for example, documents for species sensitive to one vs seven factors are 10% vs 41% likely to contain management actions. Likewise, species sensitivity is marginally related to whether climate change is considered as a threat (F(1,458)=0.33, β=0.15, p=0.07; Fig. 1a). These results indicate some prioritization of species based on potential climate threat and sensitivity, though this may be unintentional. However, overall, there is a significant gap between the sensitivity of endangered animals to climate change and the attention that climate change receives from the agencies charged with recovery of these species.

**Figure 1.**
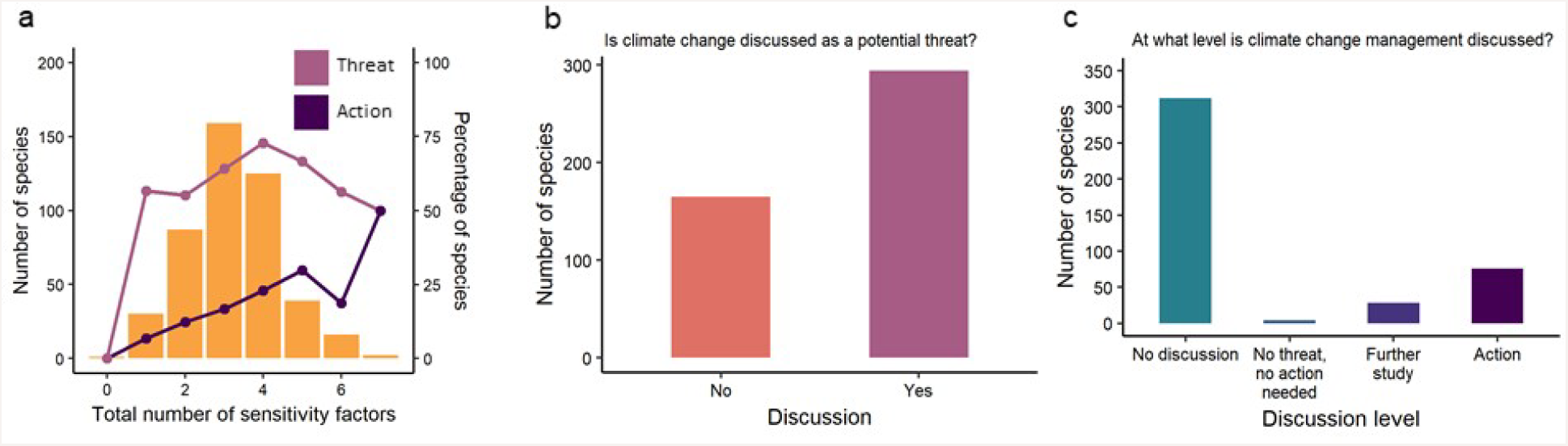
Despite sensitivity to one or more climate factors (a), US endangered animals are not often assessed for whether climate change is a potential threat (b) and most do not receive planning for management actions to address climate change impacts (c). **(a)** Species that are sensitive to more climate factors are more likely to receive management action planning (dark purple line; p<0.05) than species sensitive to fewer factors, and are marginally more likely to receive evaluation of climate change as a threat (light pink line; p=0.07). All endangered animals except one (Hawaiian goose [*Branta sandvicensis*]) are sensitive to one or more of eight climate factors (see Table 1 for description of factors). The two most sensitive species (seven factors) were a fish, the Clear Creek gambusia (*Gambusia heterochir*), and a mollusk, the shinyrayed pocketbook (*Lampsilis subangulata*). Bars represent the number of species; lines represent the proportion of species within each number of sensitivity factors. Analysis in **(a)** and **(b)** contain all endangered animals on the ESA (n=459); analysis in **(c)** excludes species for which only listing decisions exist (excluded n=39; included n=420; see text for details). Colors correspond to Fig. 2, 3, and 4.

The prevalence of sensitivity factors varied considerably. The highest proportion of species across taxa was sensitive to isolation (mean across taxa=0.71, all taxa ≥0.50), whereas the lowest proportion was sensitive to phenology (mean=0.09, all taxa ≤0.21; Fig. 2a). Hydrology and chemistry showed the highest variation in sensitivity across taxa (mean=0.60, sd=0.25, cv=0.95; mean=0.25, sd=0.22, cv=0.89, respectively); disturbance showed the least (mean=0.61, sd=0.11, cv=0.17; Fig. 2a). Of the taxa assessed, mammals were sensitive to the fewest number of factors (Fig. 2b). Amphibians, mollusks, and arthropods were sensitive to the greatest number of factors; many of these species exhibit an aquatic life cycle phase and are thus subject to hydrological and chemical sensitivities. Furthermore, mollusks and arthropods also commonly depend on obligate species relationships, although reproductive host species are not known for some mollusks. Agencies appear to be prioritizing at least some of these high-sensitivity taxa for climate change-related management. Arthropods and reptiles had the greatest proportion of species for which climate change was evaluated as a threat (80% and 75%, respectively) and management action was described (29% and 28%, respectively), whereas mollusks had the least (50% and 31%, respectively; Fig. 3a-b).

**Figure 2.**
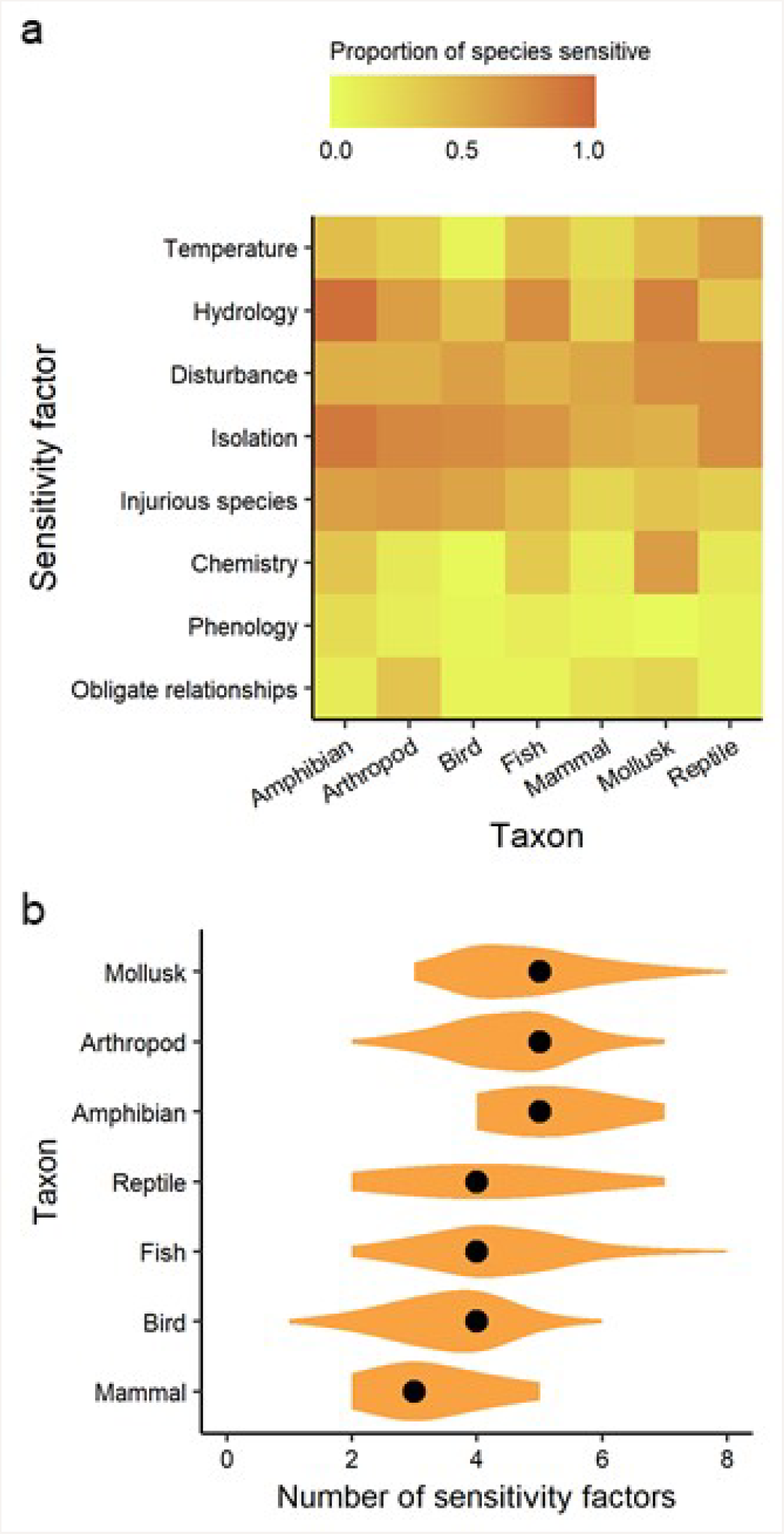
Taxa differ in sensitivity to the (a) type and (b) total number of climate factors. Analysis includes all 459 endangered species listed on the Endangered Species Act. See Table 1 for descriptions of factors, Supplementary Table 1 for the number of species in each taxa, and Supplementary Figure S1 for taxa sensitivity by factor across management agency and region.

**Figure 3.**
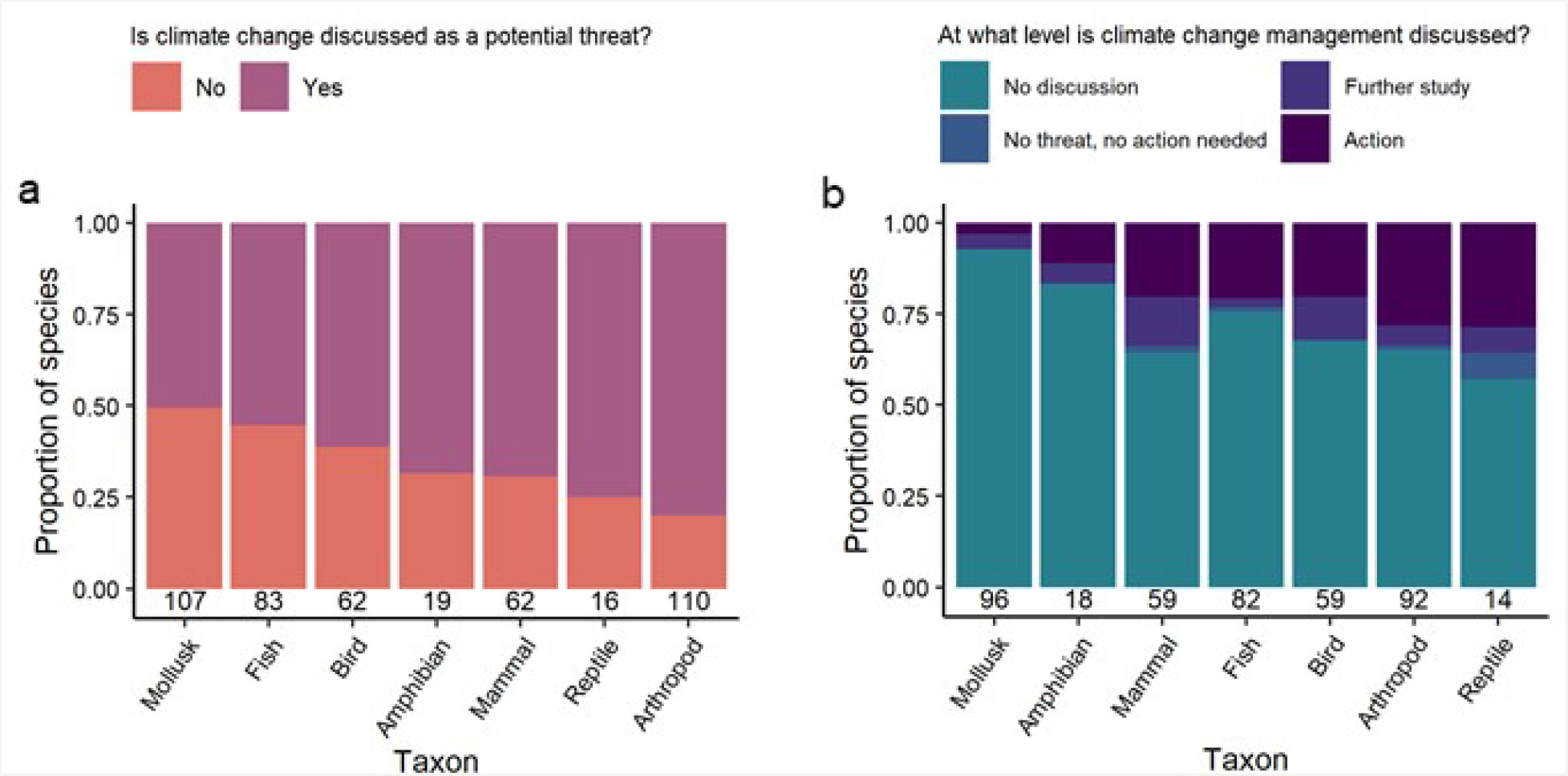
Taxonomic differences occur in whether (a) and how (b) climate change is discussed in official management documents for endangered animals. Analysis in **a** contains all 459 endangered animals listed on the Endangered Species Act; analysis in **b** excludes species for which only listing decisions exist (excluded n=39; included n=420; see text for details). The number of species in each group is shown above the x-axis.

Agencies have increasingly considered climate change as a potential threat to species in ESA documents over time, mirroring rising concern about climate change over the past few decades^26^. However, they have not yet widely translated this concern into articulated management actions to help species adapt to climate change. After the agencies first described climate change as an influence on habitat loss (listing factor 1) in 2007, the proportion of species with climate change mentioned in their ESA documents rose and thereafter stabilized at around 87% of species in 2015-2016 (Fig. 4a). In 2017-2018 however, this trend reversed, with declines in both the proportion of species where climate change was listed as a threat, and in the absolute number of newly published ESA-related documents for endangered animals. With regard to management planning, climate change was first identified as a topic for future study for the Indiana bat (*Myotis sodalis*) and Choctawhatchee beach mouse (*Peromyscus polionotus allophrys*) in 2007, and the first discussion of management action occurred in a 2008 recovery plan for the stellar sea lion (*Eumetopias jubatus*; Fig. 4b). The proportion of species with planned climate change-related action each year generally increased until peaking in 2014. Since then, discussion of action has steadily declined; of documents published in 2017, one species’ five-year review (Kaua’i cave amphipod [*Spelaeorchestia koloana*]) described a management response to climate change, and no 2018 documents mentioned actions to address climate impacts. In summary, although the number of ESA documents mentioning climate change has increased over time, most species’ documents either describe climate change as a potential problem without including any actions to specifically address the issue, or the documents do not discuss climate change at all. Across time(2007-2018), the proportion of species with planned climate change-related action has been low on average (mean=0.23, range=0.03-0.39; Fig. 4b), indicating a shortfall in planning of on-the-ground management for climate change that to date shows no sign of improving.

**Figure 4.**
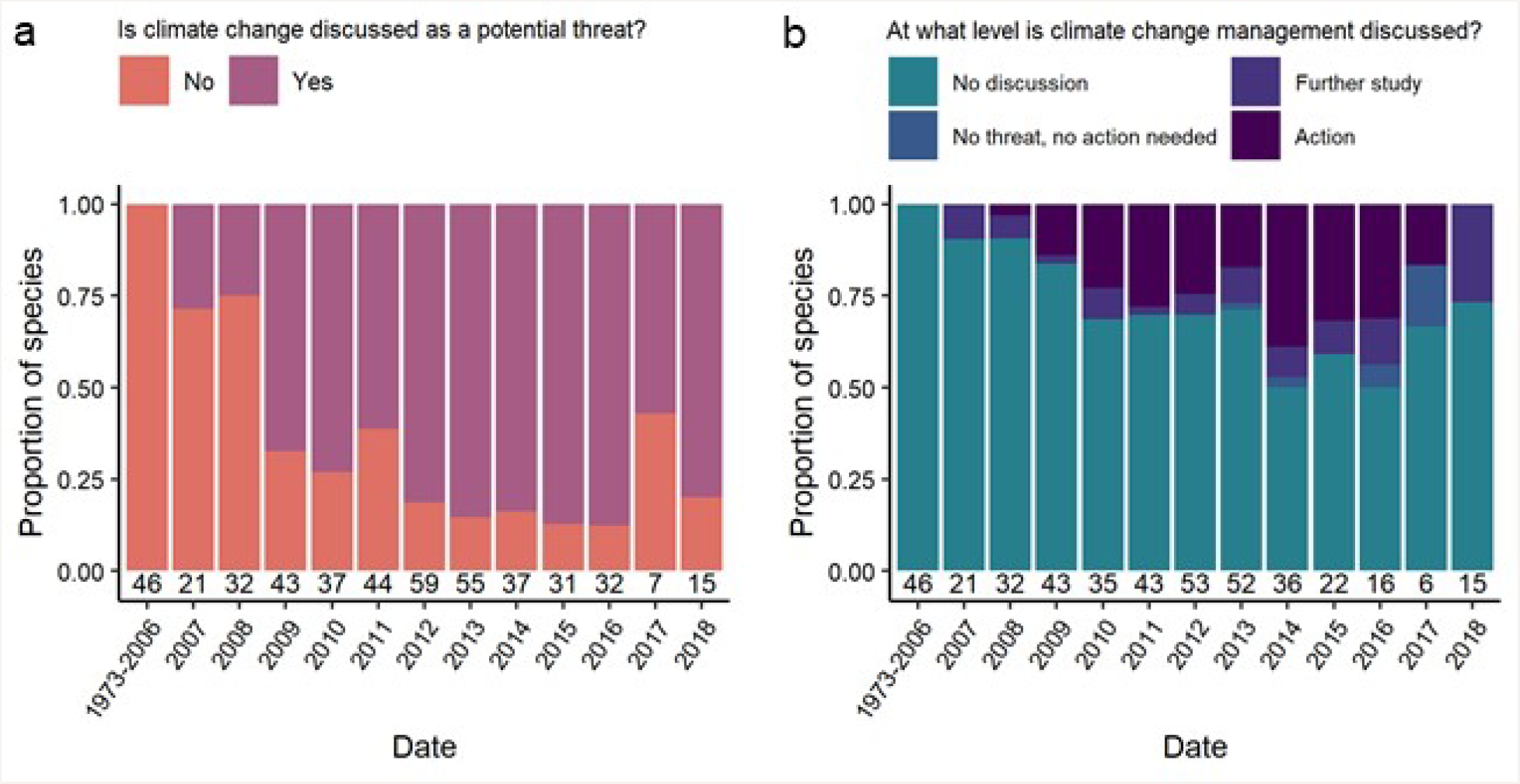
Over time US agencies have included discussions about climate change in official documents for more endangered animals, but (a) baseline assessments of climate change as a threat have increased at a substantially faster rate than (b) planning of management action. Analysis in **a** contains all 459 endangered animals listed on the Endangered Species Act; analysis in **b** excludes species for which only listing decisions exist (excluded n=39; included n=420; see text for details). The number of species in each group is shown above the x-axis.

In short, across time and taxa, management agencies are inadequately assessing climate change threats, or planning action to manage those threats, to imperiled species. In terms of baseline assessment, this inadequacy affects species regardless of their climate sensitivity, as we found no relationship between the number of sensitivity factors and the consideration of climate change as a potential threat. Agencies may be inadvertently prioritizing species for management planning based on their degree of sensitivity to climate factors, however we caution that the mere presence of management action in documents does not assure the adequacy of plans or, more importantly, the enactment of those plans^10^. Even for species with planned actions, we observed substantial variation in the content: several five-year reviews merely recommended updating recovery plans to include climate change. More robust discussions for action entailed protecting refugia (e.g., Chinook salmon [*Oncorhynchus tshawytscha*] and white abalone [*Haliotis sorenseni*] recovery plans) and diverse microsites (e.g., Karner blue butterfly [*Lycaeides melissa samuelis*] five-year review), improving connectivity (e.g., jaguar [*Panthera onca*] recovery plan), establishing additional populations for redundancy in case of stochastic climate events (e.g., Sonoran pronghorn [*Antilocapra americana sonoriensis*] recovery plan), reducing non-climate-related threats (e.g., water allocations in spikedace [*Meda fulgida*] five-year review), and designating critical habitat in areas likely to persist or become important areas in the future (e.g., tidewater goby [*Eucyclogobius newberryi*] and Bartram’s scrub-hairstreak butterfly [*Strymon acis bartrami*] critical habitat designations). Our results offer insights for how agencies, including different management jurisdictions (see Supplemental Information), might prioritize the types of climate change adaptation options to target susceptible taxa and sensitivity factors.

Three main issues may explain why the relevant US agencies have yet to address climate change threats as part of their imperiled species conservation programs. First, the politicization of climate change has caused its prioritization to shift every 4 or 8 years with changes in Presidential administration. In 2017, the Trump administration revoked many policies and commitments on climate change established by the Obama Administration, such as Executive Order 13653 on adaptation^27^ and the Paris Global Climate Agreement.^27,28^. This has disrupted progress on both mitigation and adaptation nationally and internationally^3^. Imperiled species conservation in the face of climate change urgently requires the return of a bipartisan and durable commitment to both mitigation of and adaptation to climate change. For example, legislative bodies, such as the US Congress and central governments in other countries, could integrate climate change adaptation and mitigation into law rather than leaving these important processes to more labile policies.

Second, the infrequent and inconsistent inclusion of climate change in ESA species conservation may be a consequence of chronic underfunding and imbalanced funding of species recovery. In fiscal year 2012, 62% of species recovery funding was spent on the conservation of 10% of US listed species, resulting in as little as $60 for some species (e.g., Cumberland bean mussel [*Villosa trabalis*] whose ESA documents did not mention climate change)^14,29,30^. Another analysis of yearly appropriations from 1980-2014 found that <25% of required recovery funding has been allocated annually^31^. Increased funding to the agencies responsible for species recovery, paired with a more informed allocation of resources, could help redress this problem^15,31^.

Finally, climate change itself is a formidable conservation challenge that agencies generally lack the logistical tools and capacity to address. The broad spatial and temporal scales and uncertainty of specific threats mean that agencies should pair conceptual models with mechanistic approaches to identify stressors that materialize as species threats^2,32^. Agencies would benefit from embracing the frameworks designed to enable systematic planning, implementing, and monitoring of complex conservation challenge, and integrate climate change with other threats^33,34^. Additionally, agencies should proactively seek and embrace innovative tools that enable efficient management of the 2,300+ imperiled species listed on the ESA. The assessment used in this study is one such example, offering a time-efficient method for preliminary evaluations of species sensitivity to climate change.

Our study reveals that US government agencies have yet to adequately evaluate climate change threats to endangered animals listed under the ESA and plan commensurate action. The consistency between our US results and recent findings from Australia^10,11^ suggest it is possible that many countries are similarly failing to protect imperiled species from climate change impacts. Climate change poses an ongoing and accelerating threat to many, if not most, imperiled species, and recovery will be unattainable unless a feasible process is in place to account for and ameliorate its impacts.

## Methods

We compared the climate change sensitivity of species to agency evaluation and management planning of climate change threats for ESA-listed endangered animals in the US. First, since systematic data did not exist for the climate change impacts on endangered species, we developed and conducted a trait-based, rapid assessment for evaluating climate change sensitivity. We focused the assessment on one element of species vulnerability: a species’ potential “sensitivity” to the effects of climate change. Sensitivity “refers to innate characteristics of a species or system and considers tolerance to changes in such things as temperature, precipitation, fire regimes, or other key processes”^35^. We created and answered eight yes-or-no questions based on whether the species’ habitat, ecology, physiology, or life cycle might be affected by changes in climate (Table 1). In doing so, we employed a biological approach to assessing sensitivity that considered the ecological impact to the species from the primary manifestations of climate change, including indirect impacts from effects on interacting species^24^. We derived the questions, or sensitivity factors, from factors listed in existing vulnerability assessment protocols, particularly the NatureServe Climate Change Vulnerability Index^36^ and US Forest Service’s System for Assessing Vulnerability of Species^37^. Though not exhaustive, our questions covered the main categories of species sensitivity (or sometimes categorized under adaptive capacity) in these and other assessment frameworks^38^. We were thus able to assess many of the elements of vulnerability that can be addressed via management planning, while also completing most species in 30-60 minutes. This assessment could be useful to agencies for evaluating large numbers of species while still capturing the most critical elements of potential species sensitivity.

We assessed the climate change sensitivity of all animal species listed as endangered under the ESA (as of December 31, 2018) that are found in US states, territories, and surrounding waters (n=459; see http://www.fws.gov/endangered), with the exception of those deemed likely to be extinct by agencies or which have not been observed for 20+ years and are likely extinct in the wild. We answered the assessment questions using freely-accessible species information from species listing decisions and other publicly available information published by agencies and conservation organizations about the species and its threats. We predominantly referenced the FWS’ Environmental Conservation Online System (ECOS; https://ecos.fws.gov/ecp), NMFS’ Endangered Species Conservation Directory (https://www.fisheries.noaa.gov/species-directory/threatened-endangered), and the NatureServe Explorer (http://explorer.natureserve.org). Using publicly available information enables the assessment to be used by the public or government, the latter of which requires decision data to be publicly visible^39,40^.

Consistency in measuring species sensitivity both within and between assessment tools is a recognized and ongoing challenge^41,42^. We took steps to ensure that sensitivity results were consistent within and between species in our assessment. Each species was assessed by at least two and as many as seven reviewers; species were initially reviewed by at least one of six reviewers (AC, KE, RK, SM, KT and LV) and were finally crosschecking by an expert reviewer (AD). All reviewers went through extensive training to ensure consistency in the application of the methodology (Table 1), including assessing and comparing the same species to validate and align the approach.

We also evaluated the extent to which FWS’ and NMFS’ ESA documents discussed climate change as a threat to species and included planned recovery actions to address climate change impacts. First, for all endangered animals, we recorded whether climate change was considered as a potential threat in each species’ publicly available ESA documents (listing decisions, recovery plans and outlines, critical habitat designations, and five-year reviews). We focused on the most recently published agency documents, which should reflect cumulative knowledge about the species. Then, for all endangered animals except those with only listing decisions, which are not management-oriented and thus not appropriate for evaluating management planning (n=420; excluded species n=39), we recorded what level of management action was discussed to address climate change in species recovery. We recorded the level of discussion as: “Action,” indicating that the documents articulated specific actions in response to climate change impacts; “Further study,” indicating that the agency acknowledged they require additional information before an action plan could be developed; “No threat, no action needed,” indicating that the documents discussed climate change and decided that climate change is unlikely to impede species recovery; and “No discussion,” indicating that climate change was not mentioned.

We examined patterns in sensitivity and climate change discussion by time, taxa, agency and regional jurisdiction (see Supplemental Information for latter two). We tested the relationships between the number of sensitivity factors and whether documents discussed climate change as a potential threat (yes/no) or discussed management action (by reclassifying discussion categories to create a binary variable of no action/action) using logistic regression run with the ‘stats’ package in R v.3.5.0.

## Acknowledgements

We thank Natalie Dubois and Noah Matson for valuable input and Bob Dreher, Michael Evans, Meg Evansen, Mae Lacey, Sasha Pastel, Shayna Steingard for feedback on the manuscript. Financial support for data collection was provided by University of Maryland and the Stanback Internship Program of Duke University.

## Author Contributions

A.D. and A.C. designed the study; A.D., A.C., K.E., R.K., S.M., K.T., and L.V. collected data; A.D. and J.R.B.M. analyzed data and wrote the manuscript; A.D., J.W.M., M.S., and J.R.B.M. interpreted results; J.W.M. built the web app; all authors provided critical feedback on the manuscript.

## Data Availability

Data is archived on Open Science Framework and available at https://osf.io/r9uca. A free, interactive web application containing data and results from this study is available at https://defenders-cci.org/app/ESA_climate/.

## Additional information

Supplementary information is available in the online version of the paper.

