## Supporting information for "Agency plans are inadequate to conserve US endangered species under climate change"

**Results for agencies and regional management jurisdictions**

Sensitivity of species to particular sensitivity factors generally mirrored nation-wide patterns (see main text; Fig. S1).

Analysis of discussion of climate change in management documents revealed that agencies are addressing climate change differently, and in some management jurisdictions more than others. Across agencies, documents from FWS and NMFS discussed climate change as a threat at similar proportions, for 64% (n=277) and 71% (n=17) of species under their purview, respectively (Fig. S2a; however, note the large difference in sample size). Within FWS, documents from Region 3 (Midwest) discussed climate change as a threat for 88% (n=24) of the terrestrial and aquatic species, in contrast to those from Region 5 (Northeast) which discussed climate change as a threat for only 30% (n=20) of species (Fig. S3c). With respect to planning climate change adaptation actions, differences between agencies and jurisdictions are more prominent: FWS planned actions for 17% of species (n=68) under their purview whereas NMFS planned action for 35% of species (n=8; Fig. S2b). The FWS’ Region 2 (Southwest) planned actions for 34% (n=79) of species, whereas Region 4 (Southeast), the jurisdiction with the largest number of endangered animals (n=128) included actions in documents for only 8% of species (Fig. S2d). These results offer insights into how different agencies and jurisdictions might prioritize the types of climate change adaptation options to target susceptible taxa and sensitivity factors.

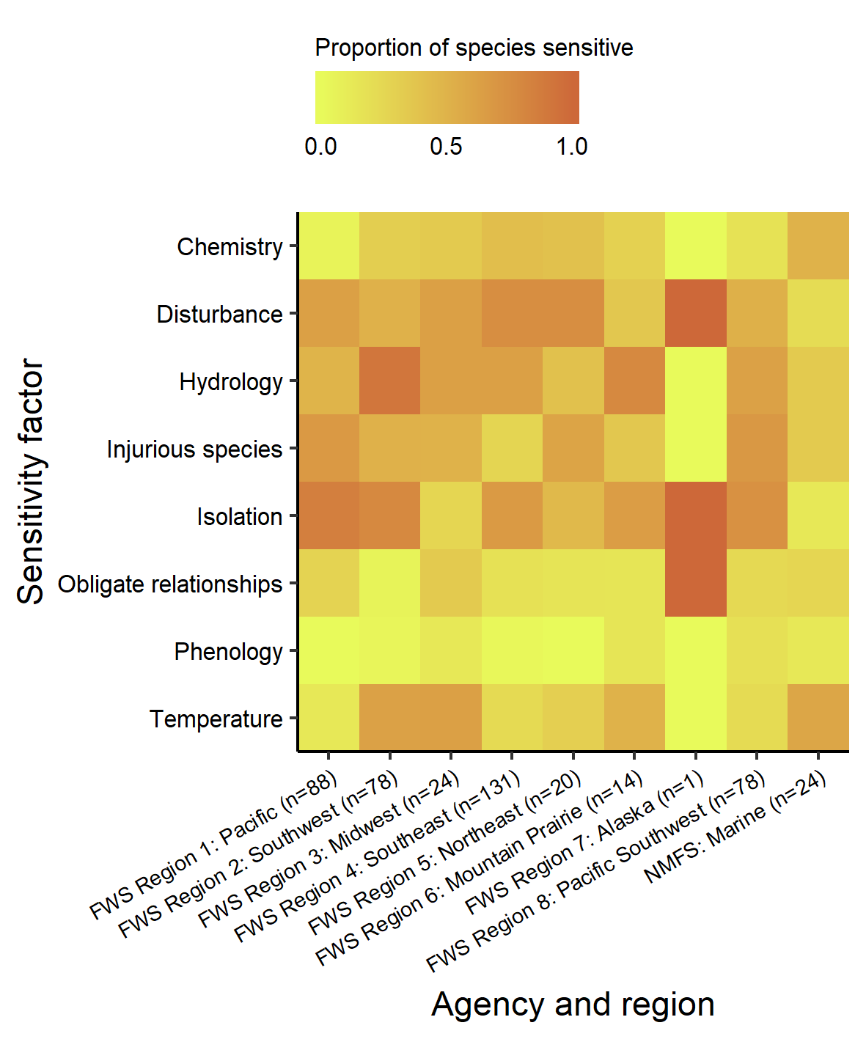

**Figure S1. The sensitivity of US endangered animals (n=459) differs by climate factor across management agency and regions.** FWS indicates US Fish and Wildlife Service and NMFS indicates US National Marine Fisheries Service. See Supplementary Table 2 for the number of species in each region and Table 1 for descriptions of climate sensitivity factors.

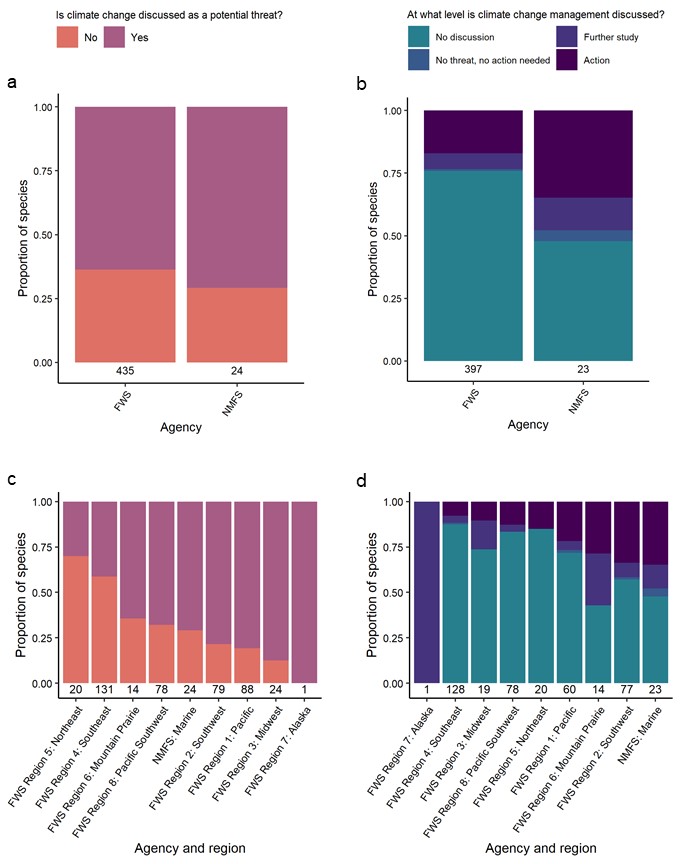

**Figure S2. Agency and regional differences occur in whether (a,c) and how (b,d) climate change is discussed in official management documents for endangered animals.** Analysis in **a** and **c** contains all 459 endangered animals listed on the Endangered Species Act; analysis in **b** and **d** excludes species for which only listing decisions exist (excluded n=39; included n=420; see text for details). The number of species in each group is shown above the x-axis. FWS indicates US Fish and Wildlife Service and NMFS indicates US National Marine Fisheries Service.

**Table 1.** Taxonomic breakdown of animals listed as endangered on the Endangered Species Act (n=459).

| **Taxon** | **Number of species** | **Percentage of species** |
| --- | --- | --- |
| Amphibian | 19 | 4 |
| Arthropod | 110 | 24 |
| Bird | 62 | 14 |
| Fish | 83 | 18 |
| Mammal | 62 | 14 |
| Mollusk | 107 | 23 |
| Reptile | 16 | 3 |

**Table 2.** Breakdown by agency and region of species listed as endangered on the Endangered Species Act (n=459). FWS indicates US Fish and Wildlife Service and NMFS indicates US National Marine Fisheries Service.

| **Agency and region** | **Number of species** | **Percentage of species** |
| --- | --- | --- |
| FWS Region 1: Pacific | 88 | 19 |
| FWS Region 2: Southwest | 79 | 17 |
| FWS Region 3: Midwest | 24 | 5 |
| FWS Region 4: Southeast | 131 | 29 |
| FWS Region 5: Northeast | 20 | 4 |
| FWS Region 6: Mountain Prairie | 14 | 3 |
| FWS Region 7: Alaska | 1 | 1 |
| FWS Region 8: Pacific Southwest | 78 | 17 |
| FWS (combined) | 235 | 91 |
| NMFS: Marine | 24 | 5 |
